# 4-state model for simulating kinetic and steady-state voltage-dependent gating of gap junctions

**DOI:** 10.1101/822007

**Authors:** M. Snipas, T. Kraujalis, K. Maciunas, L. Kraujaliene, L. Gudaitis, V. K. Verselis

## Abstract

Gap junction (GJ) channels, formed of connexin (Cx) proteins, provide a direct pathway for metabolic and electrical cell-to-cell communication. These specialized channels are not just passive conduits for the passage of ions and metabolites, but have been shown to gate robustly in response to transjunctional voltage, V_j_, the voltage difference between two coupled cells and are regulated by various chemical factors. Voltage gating of GJs may play a physiological role, particularly in excitable cells which can exhibit large transients in membrane potential during the generation of an action potential. We present a mathematical/computational model of GJ channel voltage gating to assess properties of GJ channels that takes into account contingent gating of two series hemichannels and the distribution of V_j_ across each hemichannel. From electrophysiological recordings in cell cultures transfected with Cx43 and Cx45, isoforms that are expressed in cardiac tissue, data sets were fit simultaneously using global optimization. The results showed that the model is capable of describing both steady-state and kinetic properties of homotypic and heterotypic GJ channels composed of these connexins. Moreover, mathematical analyses showed that the model can be simplified to a reversible two-state system and solved analytically, using a rapid equilibrium assumption. Given that excitable cells are arranged in interconnected networks, the equilibrium assumption allows for a substantial reduction in computation time, which is useful in simulations of large clusters of coupled cells. Overall, this model can serve not just as a modeling tool, but also to provide a means of testing GJ channel gating behavior.

**Significance:** Gap junction (GJ) channels gate in response to transjunctional voltage which provides the capacity for dynamic regulation of intercellular coupling. Kinetic properties of GJs in modeling studies have been infrequently addressed and we present a computational model of voltage gating that can account for both kinetic and steady-state changes in junctional conductance, g_j_. Although GJs possess two gating mechanisms, our analysis indicates that changes in g_j_ for each voltage polarity can be adequately described by a kinetic scheme describing a single mechanism in each of the hemichannels, suggesting functional dominance of one mechanism over a substantial voltage range. This property allowed for model simplification that can be applied for efficient simulation of sizeable cell clusters and analyses of electrophysiological data.

## Introduction

Gap junction (GJ) channels are intercellular channels that mediate the direct transfer of ions, metabolites and small signalling molecules between cells in virtually all tissues in the body. In vertebrates, GJ channels are formed of connexin (Cx) protein subunits and in humans there are 21 different Cx isoforms that show tissue-specific, but overlapping patterns of expression [1, 2]. It is well established that GJ channels are sensitive to transjunctional voltage, V_j_, the voltage difference between cells [3, 4]. Sensitivity to V_j_ differs considerably among Cx isoforms as do biophysical properties such as unitary conductance and modulation by factors such as pH, Ca^2+^ and chemical agents. As advances in imaging are making recordings from cellular networks more feasible, of interest is to have models that can incorporate voltage gating of GJs, particularly in excitable tissues such as in the nervous system and the heart, where gating can play a role. In this study, we present a mathematical/computational model that can adequately describe the kinetic and steady state changes in junctional conductance, g_j_, and its dependence on V_j_. We suggest that this model could be efficiently applied for analyses of electrophysiological recordings and simulations of electrically coupled clusters of cells.

The first mathematical model of voltage gating in GJs was published in a series of papers by Bennett, Harris and Spray [5–7]. Working on pairs of amphibian blastomeres, g_j_ was found to be maximum at V_j_=0 and decrease symmetrically for both polarities of V_j_. A Boltzmann function was applied to each polarity of V_j_ to describe the steady-state g_j_-V_j_ relationship [5]. This type of analysis became a standard way to quantitate sensitivity of GJs to V_j_. The same authors also presented a mathematical model to describe the kinetic properties of GJ voltage gating [6]. The changes in g_j_ over time were evaluated using a two-state (so-called independent gating model) and a three-state (so-called contingent gating model), which described opening and closing of GJ channels containing two mirror symmetrical gates in series. These modelling studies were performed well before the identification of Cx genes and a detailed knowledge of the varied properties of GJ channels at macroscopic and single channel levels.

Over the years, biophysical, molecular and structural studies have established that GJ channels are formed by the docking of two hemichannels, which can be composed of a single or mixed combinations of Cxs, and that it is the individual hemichannels that gate in response to V_j_ to give rise to the overall behaviour of a GJ channel. The sensitivity of GJ channels to V_j_ and not the absolute membrane voltages of the two coupled cells indicates that the voltage-sensing elements detect the local electric field within the pore. This property means that the gating of one hemichannel will affect the other through alterations in the field, which was termed contingent gating by Harris et al, 1981. The wealth of biophysical data generated at macroscopic and single channel levels has prompted the generation of a series of gating models of increasing complexity. Vogel and Weingart (1998) presented a four-state model, which described the behaviour of the two apposed hemichannels, each capable of transiting between high and low conductance states, each exhibiting conductances that varied exponentially with voltage [8]. This model was used to simulate a linear array of cells coupled by Cx43-like GJ channels to assess changes in g_j_ that could occur during impulse propagation [9]. An identical four-state scheme, but using the Boltzmann gating parameters described in Spray et al (1981) [5] was applied in another modelling study [10]. The authors derived mathematical formulas for evaluation of steady-state g_j_-V_j_ relationships and validated mathematical modelling results with data obtained from electrophysiological recordings in cells expressing homotypic and heterotypic GJs. However, the kinetics of GJ channel gating was not addressed. Building on this last model, Bukauskas and colleagues developed a series of models starting with a stochastic four state model, S4SM, [11]. A subsequent 16-state (S16SM) model [12] incorporated the existence of two gating mechanisms in each hemichannel, termed fast or V_j_ gating and slow or loop gating [3]. This model was extended to a 36-state (36SM) model to include a second closed state in each hemichannel, termed the deep-closed state, associated with the slow or loop gating mechanism [13]. 36SM was applied in the modelling of neuronal networks [14]. S4SM, S16SM and 36SM all used a theoretical parameter, P_t_, to describe gating kinetics and to calibrate timescales of electrophysiological and simulation experiments. Global optimization methods were used to estimate other model parameters. However, this theoretical parameter, Pt, did not reflect any biophysical property of a GJ channel and did not depend on V_j_, and thus rendered S4SM and S16SM inadequate for describing gating kinetics. The 36-state model was more effective in that respect, perhaps by the increased number of adjustable parameters and system states, but came at a high computation cost, making it difficult to apply to simulations of large clusters of cells, such as cardiac or nervous tissue. Also, the high number of model parameters in 36SM, which could exceed 20 in a heterotypic GJ channel, made it very difficult to find a reliable, unique solution.

Our motivation here was to create a simple and computationally efficient mathematical model that could adequately describe both kinetic and steady-state properties of voltage-dependent gating in GJ channels. Building on the aforementioned research by us and other groups, we present here a model which uses a 4-state kinetic scheme to describe opening and closing of two apposing gates in series, one in each hemichannel as previously described in [8, 10, 11]. For a description of gating kinetics, we use the assumption, as did Harris et al., 1981 [6], that the free energy difference between system states depends linearly on V_j_. However, in our model, we account for the distribution of V_j_ across each hemichannel as in Paulauskas et al., 2009 [11]. To demonstrate the validity of our model, we performed model fitting of electrophysiological recordings obtained in cells expressing homotypic Cx43 and Cx45 and heterotypic Cx45/Cx43EGFP GJs. For parameter estimation we applied global optimization methods as in our previous studies [13], just in a more systematic way. Model fitting results showed good correspondence with both kinetic and steady-state data. In addition, mathematical analyses showed that the current model can be approximated by a reversible two-state system and solved analytically using a rapid equilibrium assumption [15], which is often applied in modelling enzyme kinetics [16]. This model property allows for a substantial reduction in computation time, and could be efficiently applied when simulating large clusters of cells.

## Methods

### Cell lines and culture conditions

Experiments were performed in HeLa cells, transfected with Cx45, and Novikoff cells endogenously expressing Cx43. Cell cultures were grown in DMEM supplemented with 10% fetal calf serum, 100 mg per ml streptomycin and 100 units per ml penicillin, and maintained in a CO_2_ incubator (37°C and 5% CO_2_).

### Electrophysiological recordings

Electrophysiological recordings were performed in a modified Krebs–Ringer solution containing the following (in mM): 140 NaCl, 4 KCl, 2 CaCl_2_, 1 MgCl_2_, 2 CsCl, 1 BaCl_2_, 5 glucose, 2 pyruvate, 5 HEPES, pH 7.4. Recording pipettes (3-5 MΩ) were filled with standard pipette solution containing the following (in mM): 130 CsCl, 10 NaAsp, 1 MgCl_2_, 0.26 CaCl_2_, 2 EGTA and 5 HEPES (pH 7.3).

Junctional conductance was measured in selected cell pairs using a dual whole-cell patch-clamp system. Each cell within a pair was clamped with a separate patch-clamp amplifier (EPC-7plus, HEKA). V_j_ was induced by stepping the voltage in one cell while keeping it constant in the other. Junctional current (I_j_) was measured as the change in the current of a neighboring cell and g_j_ was estimated from the relationship g_j_ = −I_j_/V_j_.

Signals were acquired and analyzed using an analog-to-digital converter (National Instruments, Austin, TX) and custom-made software [17].

### Computational modelling

Numerical solution of the computational model was implemented in MATLAB (see Supporting Material for program codes). Model fitting and parameter estimation were performed using MATLAB’s Global Optimization Toolbox.

## Model description

### Model states

The model describes a gap junction (GJ) channel containing two apposing hemichannels docked in a head-to-head fashion. Each hemichannel contains a voltage sensitive gate, which transits between an *open* (*O*) state and a *closed* (*C*) state (Fig. 1A), where the closed state typically is characterized by a residual conductance due to incomplete occlusion of the pore. The *O*↔*C* gating transitions depend on the voltage across each hemichannel (Fig. 1B).

**Figure 1.**
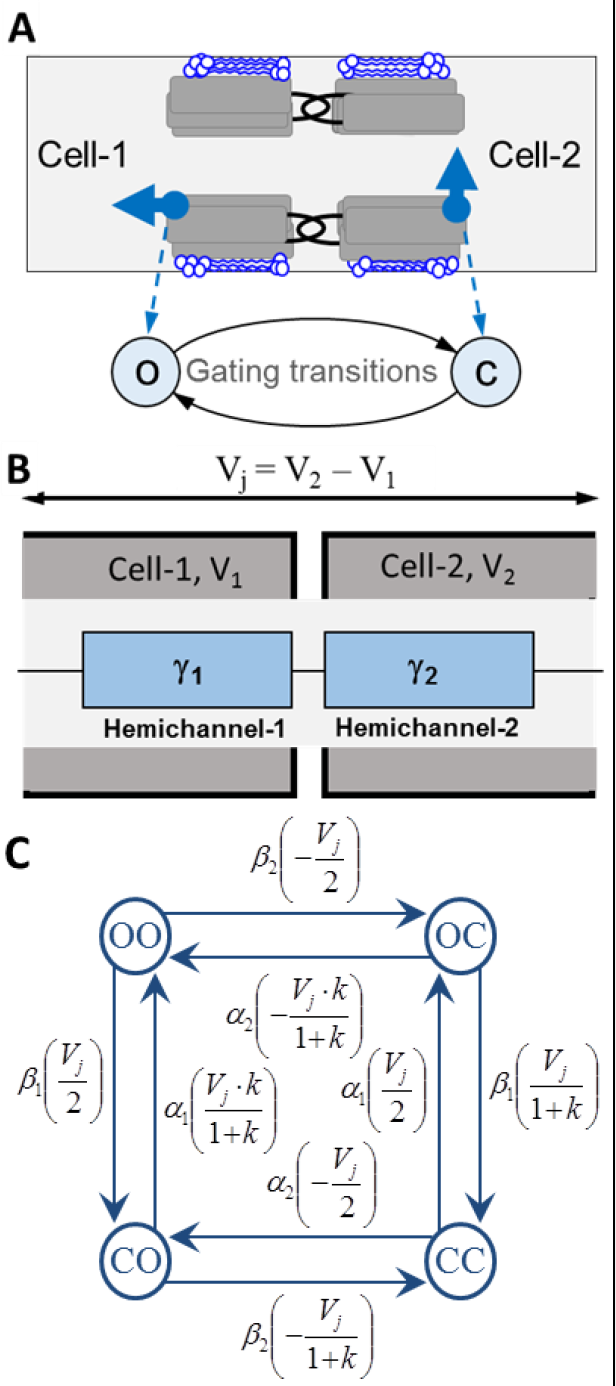
Schematics of gap junction (GJ) channel and 4-state model (4SM). (**A**) Depiction of a GJ channel, connecting two adjacent cells. Each apposing hemichannel contains a gate (blue arrows), which can transit between an open (*o*) and closed (*c*) states. (**B**) A schematic of two hemichannels with associated conductances (γ) arranged in series. (**C**) Kinetic scheme of 4SM for a homotypic GJ channel. Transition rates, α, β, depend on the transjunctional voltage (V_j_) that is distributed across each hemichannel (see Table 1).

**Table 1.** Voltage distribution across non-rectifying heterotypic and homotypic GJ channels in 4SM

|  | Heterotypic GJ channel |  | Homotypic GJ channel |  |
| --- | --- | --- | --- | --- |
| state | Hemichannel-1 | Hemichannel-2 | Hemichannel-1 | Hemichannel-2 |
| *OO* | $\frac{V_j \cdot \gamma_{2;o}}{\gamma_{1;o} + \gamma_{2;o}}$ | $-\frac{V_j \cdot \gamma_{1;o}}{\gamma_{1;o} + \gamma_{2;o}}$ | $\frac{V_j}{2}$ | $-\frac{V_j}{2}$ |
| *OC* | $\frac{V_j \cdot \gamma_{2;c}}{\gamma_{1;o} + \gamma_{2;c}}$ | $-\frac{V_j \cdot \gamma_{1;o}}{\gamma_{1;o} + \gamma_{2;c}}$ | $\frac{V_j}{1+k}$ | $-\frac{V_j \cdot k}{1+k}$ |
| *CO* | $\frac{V_j \cdot \gamma_{2;o}}{\gamma_{1;c} + \gamma_{2;o}}$ | $-\frac{V_j \cdot \gamma_{2;o}}{\gamma_{1;c} + \gamma_{2;o}}$ | $\frac{V_j \cdot k}{1+k}$ | $-\frac{V_j}{1+k}$ |
| *CC* | $\frac{V_j \cdot \gamma_{2;c}}{\gamma_{1;c} + \gamma_{2;c}}$ | $-\frac{V_j \cdot \gamma_{2;c}}{\gamma_{1;c} + \gamma_{2;c}}$ | $\frac{V_j}{2}$ | $-\frac{V_j}{2}$ |

Considering that either hemichannel in a GJ channel can transit between open or closed states, a GJ channel exhibits four different states:

*OO* – hemichannel-1 open, hemichannel-2 open;
*OC* – hemichannel-1 open, hemichannel-2 closed;
*CO* – hemichannel-1 closed, hemichannel-2 open;
*CC* – hemichannel-1 closed, hemichannel-2 closed.

### The voltage distribution across hemichannels

Because hemichannels gate by sensing the local electric field in the pore, the voltage distribution across each hemichannel is determined by the states of the two hemichannels and the conductances associated with each state. Thus, broadly, GJ channels exhibit contingent gating, as introduced in [6], in which gating of each hemichannel is contingent on the state of the other. Referring to two cells, termed cell 1 and cell 2, the relevant voltage is the transjunctional voltage, V_j_, which is defined as the voltage difference between the cells, i.e. V_j_=V_2_−V_1_. Because the hemichannels are docked in a head-to-head fashion, the voltage sensing elements in each hemichannel are oriented in opposite directions and, thus, sense V_j_s that are, in essence, opposite in polarity. In our model, the polarity of the voltage drop across the hemichannel in cell 1 is defined as the same polarity as the applied V_j_, whereas the polarity of the voltage drop across the hemichannel in cell 2 is of opposite sign.

Because unitary conductance is usually constant with voltage in homotypic GJs [4], we only evaluated rectification of unitary conductance in some of our numerical experiments considering heterotypic GJ channels. Assuming a constant unitary conductance, the magnitude of the voltage drop across each hemichannel can be estimated from a simple equation describing a voltage divider circuit, as presented in Table 1. For homotypic GJs, the calculation can be simplified given that the apposing hemichannels have the same open and closed state conductances, i.e., *γ*_1;*o*_ = *γ*_2;*o*_ = *γ*_*o*_ and *γ*_1;*c*_ = *γ*_2;*c*_ = *γ*_*c*_. For example, if we denote the ratio between closed and open hemichannel conductances as *k* = *γ_c_*/*γ_o_* (0 ≤ *k* ≤ 1), the V_j_ distribution can be expressed as presented in the 3^rd^ and 4^th^ columns of Table 1:

### Rectification of unitary conductance

Rectification of unitary conductance was evaluated using the same method as originally proposed in [8], and later applied in our modelling studies [13, 18]. That is, we presumed that the unitary conductance (γ) of an open or a closed hemichannel is a function of voltage (V) according to:

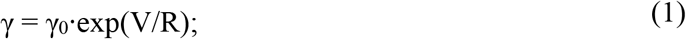

where γ_0_ denotes the unitary conductance at V_j_=0 mV and R is a coefficient that defines the steepness of rectification. In this case, the voltages across the first and the second hemichannels, V_1_ and V_2_ respectively, must satisfy the following system of nonlinear equations at each of system states:

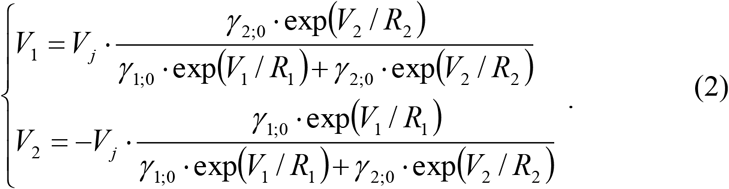

Here indices 1 and 2 in the subscripts of the variables γ_0_ and R denote the unitary conductances of the two hemichannels at V_j_=0 mV and their rectification coefficients, respectively.

The nonlinear system of Eq. 2 does not have an explicit closed form solution and must be solved numerically. Similarly to our previous studies [12, 18], we found out that a fixed-point iterative method can be successfully applied to evaluate the voltage distribution across the gates. It has a high speed of convergence and can provide a solution of sufficient precision (<0.0001) in just a few iterations.

### Transition rates of hemichannel gating

The rates of *o*↔*c* gating transitions were estimated with the assumption that the free energy difference between system states depends linearly on the voltage across each hemichannel [6]. In such a case, opening (*C*→*O*) and closing (*O*→*C*) rate constants, and respectively, are exponential functions of voltages across hemichannels (V):

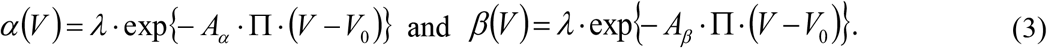

Here A_α_ and A_β_ describe the voltage sensitivities of the respective transition rates; Π describes hemichannel gating polarity (Π = −1, if hemichannel tends to close at negative voltage, and Π = 1 otherwise); V_0_ is voltage, at which hemichannel opening and closing transition rates are equal (i.e., at equilibrium, half of hemichannels are open at V_0_); λ is the hemichannel opening and closing rate at V_0_.

Based on the aforementioned assumptions, the ratios of channels in each of 4 states at time *t*, or alternatively, the probabilities of channels states, *oo*(*t*), *oc*(*t*), *co*(*t*) and *cc*(*t*) can be described by the following system of ordinary differential equations (ODEs):

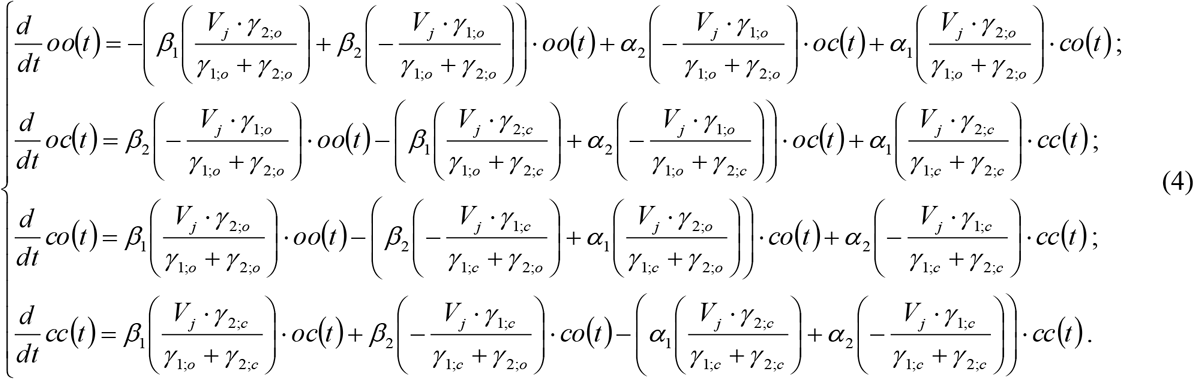

State variables *oo*(*t*), *oc*(*t*), *co*(*t*) and *cc*(*t*) denote probabilities of channel states and thus, they conform to the following constraints: 0 ≤ *oo*(*t*), *oc*(*t*), *co*(*t*), *cc*(*t*) ≤ 1. In addition, because insertion of new channels or degradation of channels over time is not considered here, the following conservation holds at all times: *oo*(*t*)+*oc*(*t*)+*co*(*t*)+*cc*(*t*)=1.

Eq. 4 describes the kinetics of the 4-state model (4SM), which is applicable for both homotypic and heterotypic GJ channels. Fig. 1C shows the kinetic scheme for a homotypic GJ channel, for which the V_j_ distributions across hemichannels can be estimated as presented in Table 1.

Once *oo*(*t*), *oc*(*t*), *co*(*t*) and *cc*(*t*) are estimated, overall junctional conductance (g_j_) at time *t* can be estimated as an averaged conductance of each channel state, weighted according to its probability:

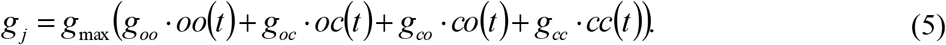

Here, *g_max_* denotes maximum conductance, which is achieved when *oo*(*t*)=1 (that is, all channels are in the open state *OO*); *g_oo_*, *g_oc_*, *g_co_* and *g_cc_* denote the junctional conductances at each system state, which can be estimated from the rule of resistors in series:

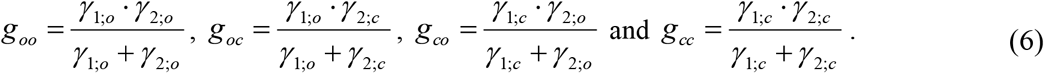

For a homotypic GJ channel, these expressions can be simplified as follows:

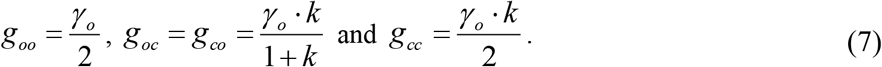

here, as in Table 1, *k* denotes the ratio of the unitary conductances of closed and open hemichannels, *k* = γ_*c*_/γ_*o*_.

## Results

### Model fitting of 4SM to evaluate junctional conductance of homotypic GJ channels

We fit electrophysiological data obtained in HeLa cells expressing Cx45 using 4SM. Fig. 2 shows model fitting results for averaged reductions in g_j_ in response to various negative V_j_ steps. Average g_j_ values and standard errors in Fig. 2 were obtained from at least 5 different experiments. Model parameters of homotypic Cx45 GJs were estimated using global optimization methods. Optimization was performed to provide the best fit to all the different V_j_ steps simultaneously. We assumed that the apposing hemichannels in any homotypic GJ channel exhibit identical V_j_ gating properties. In addition, we assigned a negative gating polarity to Cx45 hemichannels, as reported in [19]. Thus, a Cx45 hemichannel in cell 1 would tend to close when the applied V_j_ is negative.

**Figure 2.**
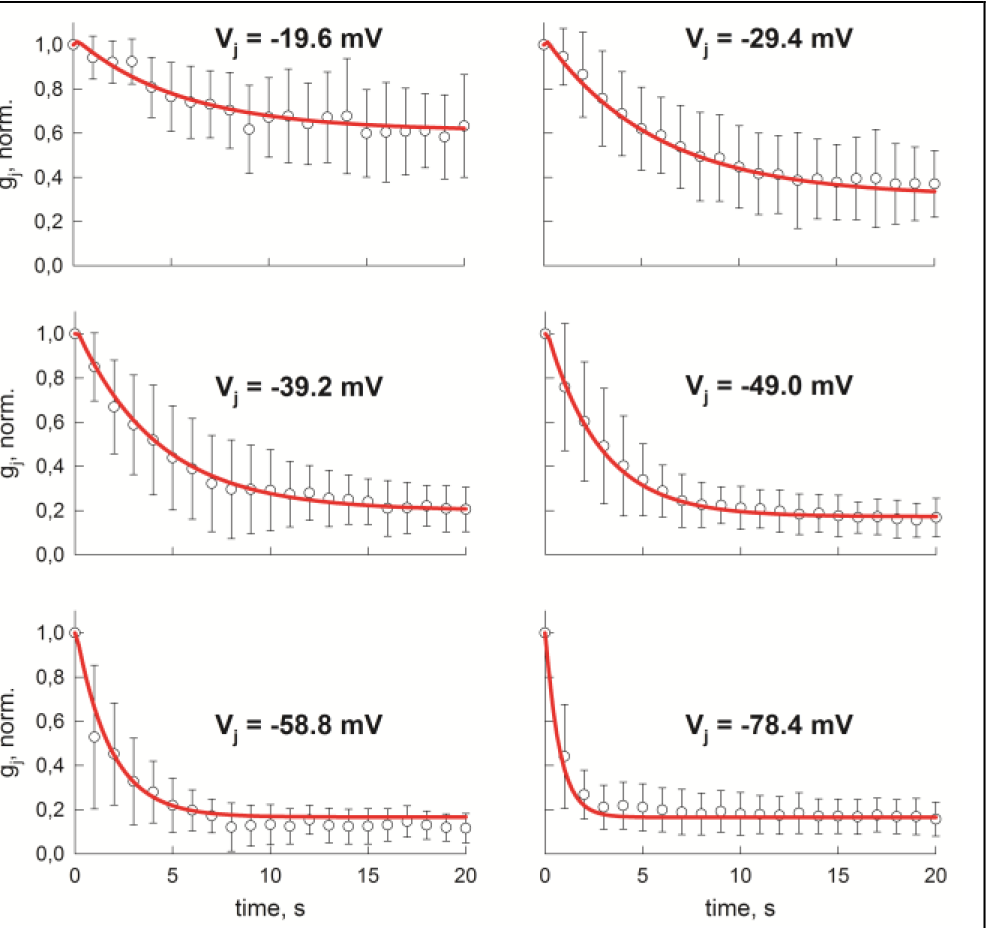
Model fitting of 4SM to reductions in gj in response to negative V_j_ steps. Open circles show averaged gj time courses recorded in HeLa cells expressing Cx45. Normalized average values of g_j_ and standard errors were estimated from at least 5 experiments. Red lines show g_j_ time courses evaluated using 4SM. Model parameters were estimated to fit all experimental data simultaneously.

Fig. 2 demonstrates that 4SM provides a good fit (red lines) to the changes in g_j_ over time for all applied V_j_ steps. Global optimization provided the following estimates of model parameters: λ=0.1415 s^−1^, A_α_=0.1264 mV^−1^, A_β_=0.0920 mV^−1^, V_0_ = −14.35 mV and γ_*o*_/γ_*c*_ = 0.1665.

To further test the model, we examined how well this same set of model parameters described the steady-state g_j_-V_j_ relationship obtained from a previously published data set for Cx45 GJs [20]. The results in Fig. 3 show that the 4SM (solid black line in Fig. 3A) provides a good fit to the experimental data (white circles) and provides an independent validation of the model because data in Fig. 3 were not used in obtaining model parameters.

**Figure 3.**
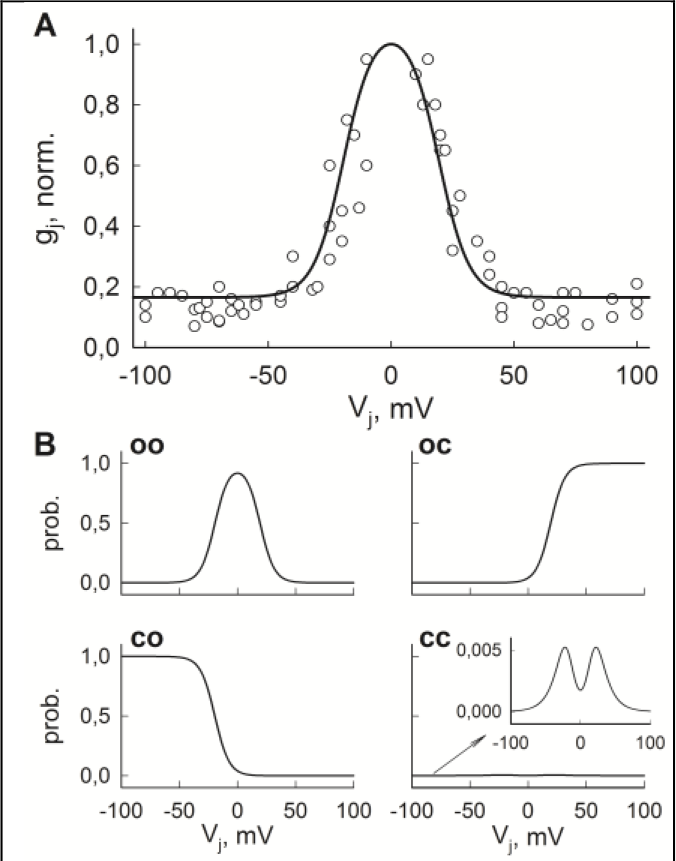
Model fitting of 4SM to steady-state g_j_-V_j_ relationship of Cx45 GJ channels and steady-state probabilities. (A) Open circles show normalized steady-state g_j_-V_j_ values, recorded in HeLa cells, expressing Cx45. The black line denotes g_j_-V_j_ relationship obtained from 4SM that was fit to the kinetic data set shown in Fig. 2. (B) Steady-state probabilities of all the states in 4SM. Here, we used the same set of model parameters as in Fig. 2, which were estimated independently from data presented in A.

Fig. 3B shows estimates of steady-state probabilities for all four model states, using the same set of model parameters. According to the model, at V_j_=0 mV about 90 percent of homotypic Cx45 GJ channels reside in an open (*OO*) state, and the macroscopic residual conductances for either polarity of V_j_ are largely determined by the states with a single hemichannel closed (*OC* and *CO*). The probability of residing in the state with both hemichannels closed (*CC*) is predicted to be rather uncommon, reaching a peak probability of ~0.006 at around V_j_=±20 mV (see insert in Fig. 3B). Our model fitting estimates a unitary conductance of ~3 pS for the *CC* state of a homotypic Cx45 GJ, which would be difficult to measure experimentally due to the noise inherent in dual whole-cell recordings.

We also performed model fitting for homotypic Cx43 GJ channels. In this case, experimental data was obtained using V_j_ ramp protocols (see Fig 4A), which resulted in hysteresis of the g_j_-V_j_ relationship when g_j_ was calculated using increasing and decreasing segments of the V_j_ ramps. The electrophysiological recordings were obtained in Novikoff cells that endogenously express Cx43. Model fitting was performed using data averaged from 5 applied ramps in a single cell pair. Fig. 4A-B shows that 4SM provides a good fit (solid red curves), except for, perhaps, at the end of recovery in which the experimental g_j_ values did not fully recover.

**Figure 4.**
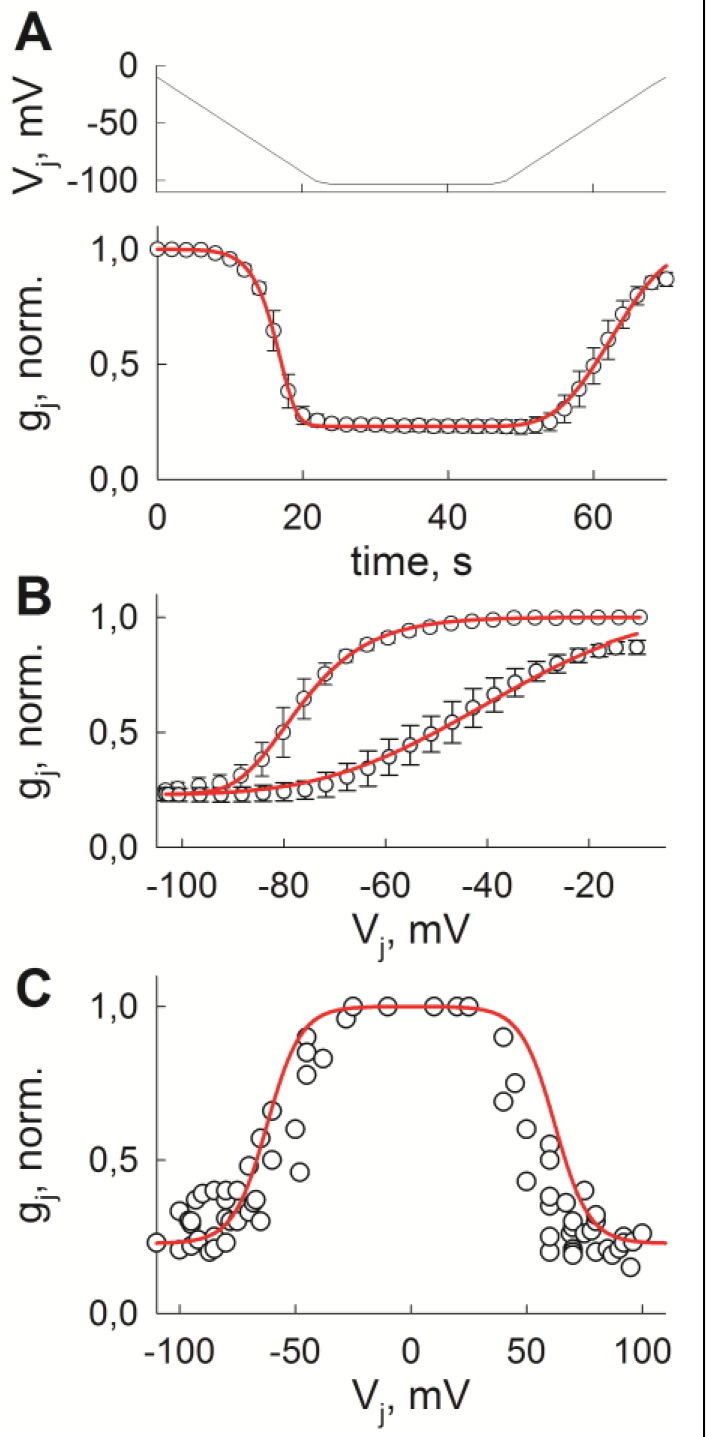
Model fitting using 4SM can describe g_j_ kinetics, including hysteresis, in response to increasing and decreasing V_j_ ramps. **(A)** Time course of g_j_ decrease and recovery (lower panel) in response to V_j_ ramp protocol (upper panel). Average g_j_ values and standard errors (open circles and error bars) were obtained from 5 different V_j_ ramp protocols in a single cell pair. Electrophysiological recordings were obtained in Novikoff cells, which endogenously express Cx43. **(B)** The same data as in **A**, just plotted to show hysteresis of the g_j_-V_j_ relationships. In both cases, solid red curve was fitted using 4SM. **(C)** Steady-state g_j_-V_j_ relationship in Cx43 GJs. Open circles show experimentally observed g_j_-V_j_ values recorded at the end of sufficiently long V_j_ steps. Solid red curve was obtained using 4SM with the same model parameters as in **A** and **B**, which were fitted independently from data in **C**.

The model parameters that were obtained were as follows: λ=0.1522 s^−1^, A_α_=0.032 mV^−1^, A_β_=0.215 mV^−1^, V_0_ = −34.24 mV and γ_*o*_/γ_*c*_ = 0.257. Gating polarity was assigned to be negative, as previously reported in [21]. Again, for independent validation, the steady-state g_j_-V_j_ relationship generated using the same set of parameters (Fig. 4 C; red solid line), was superimposed on the steady-state g_j_-V_j_ values (white circles) obtained from a separate experimental set [22]. As for Cx45 GJs, the simulation results and experimental data are in good agreement for Cx43 GJs. Overall, the results from Cx45 and Cx43 model fitting indicate that 4SM can adequately describe both kinetic and steady-state V_j_ gating properties of homotypic GJ channels.

### Model simplification using the rapid equilibrium assumption

Analysis of 4SM modeling results using Cx45 and Cx43 gating parameters revealed that different pairs of variables *oo*(*t*), *oc*(*t*), *co*(*t*) and *cc*(*t*) reach equilibrium ratios at different timescales in response to a V_j_ step. For example, at negative applied V_j_s, the equilibrium ratio between *oo*(*t*) and *oc*(*t*), as well as between *co*(*t*) and *cc*(*t*), was achieved very rapidly, while later transitions from states *OO* and *OC* to *CO* and *CC* occur at much slower timescale (see Fig. 5A). These kinetics are the result of much higher *OO*↔*OC* and *CO*↔*CC* transition rates at negative V_j_s and allowed us to simplify the model using a rapid equilibrium assumption. We assumed that pools of channels residing in states *OO* and *OC*, as well as in *CO* and *CC*, are in equilibrium during simulation with negative V_j_ steps. This assumption led to a reduced model with only two variables: 1) *o*_1_(*t*)=*oo*(*t*)+*oc*(*t*), which denotes the proportion of channels with hemichannel-1 open, and 2) *c*_1_(*t*)=*co*(*t*)+*cc*(*t*), which denotes the proportion of channels with hemichannel-1 closed. Our analysis showed that this two state model can be described by a reversible kinetic scheme 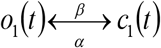 with the following closing (β: *o*_1_→*c*_1_) and opening (α: *c*_1_→*o*_1_) transition rates (full model derivation is presented in Supporting Material):

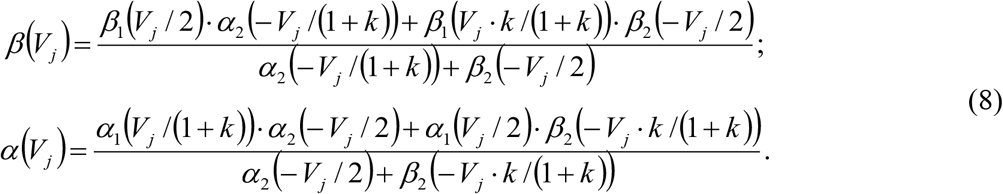

**Figure 5.**
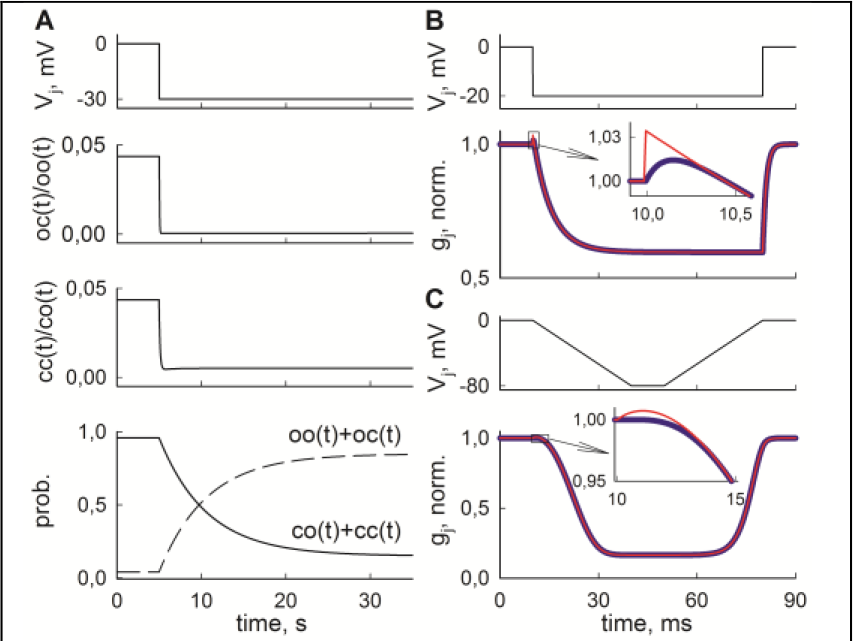
Model simplification, based on a rapid equilibrium assumption (REA). **(A)** Kinetics of 4SM variables show a clear separation of timescales in response to a negative V_j_ step (upper panel). Rapid equilibrium ratios are achieved between pairs of variables (middle panels), which denote that hemichannel-1 is open (*oo*(*t*) and *oc*(*t*)) or closed (*co*(*t*) and *cc*(*t*)). On a longer timescale, channels with hemichannel-1 open transit to states with hemichannel-1 closed (lower panel). **(B)** and **(C)** The comparison of modeling results using the full 4SM and the simplified model (thick blue and thin red curves in g_j_ time courses, respectively), which is based on REA approximation. The curves basically overlap and small differences between the outputs of two models can only be seen at the beginning of V_j_ step or ramp protocols (see inserts). In all cases, we used the same gating parameters, which were fit to Cx45 GJs.

Here transition rates on the right sides of the equations are the same as in the full model (see Fig. 1C).

The solution of a two-state reversible system is well known with kinetics that can be described by an exponential relaxation to a steady state. In this case, the time course of *o*_1_(*t*) and *c*_1_(*t*) after a negative V_j_ step of duration *t* can be expressed as follows:

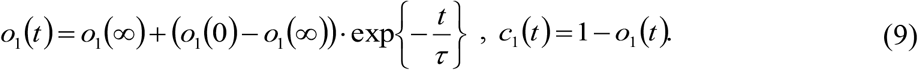

Here *o*_1_(0) denotes an initial value of *o*_1_ at the beginning of negative V_j_ step, *o*_1_(∞) is the steady-state value given by 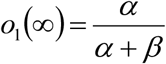, and τ is a time constant given by 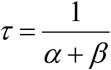. Then, state probabilities *oo*(*t*), *oc*(*t*), *co*(*t*) and *cc*(*t*) can be estimated as follows:

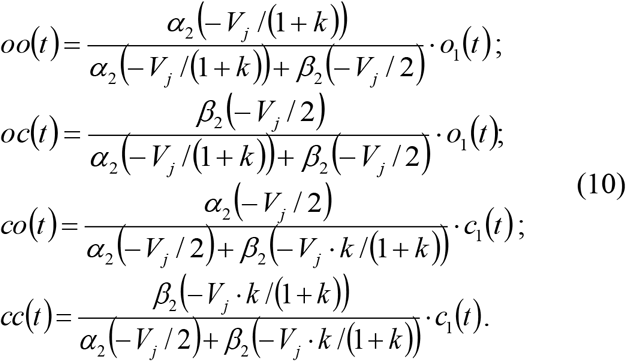

A similar analysis can be performed for positive V_j_ steps. In that case, a rapid equilibrium ratio is achieved between pairs of states in which hemichannel-2 is open (*OO* and *CO*) or closed (*OC* and *CC*). Thus, the kinetics between pools of states *o*_2_(*t*)=*oo*(*t*)+*co*(*t*) and *c*_2_=*oc*(*t*)+*cc*(*t*) can also be described by a two-state reversible system. Estimation of transition rates for V_j_>0 is presented in the Supporting Material.

### Application of the simplified model

An important aspect of a simplified model is a very significant reduction in computation cost. That is, the full 4SM requires solving a system of four ODEs (2), e.g., by calculating an exponential matrix. The simplified model allowed us to replace these computationally costly operations with analytical solutions, provided by Eqs. 9 and 10. We compared simulated g_j_ time courses in response to various V_j_ step and ramp protocols using both the full 4-state model and the simplified model, derived from rapid equilibrium assumption (REA). We used the same Cx45 and Cx43 V_j_ gating parameters, obtained from model fitting results (see Figs. 2-4).

Initially, the REA approximation was unable to adequately describe recovery of g_j_ at V_j_=0 mV, following a positive or negative V_j_ step. This problem was solved by observing that left and right hemichannels act independently at V_j_=0 mV, and can be described as two separate 2-state processes with reversible gating kinetics. A full mathematical derivation of this property of 4SM, together with a program code for a simplified model is provided in Supporting Material.

Fig. 5B shows two examples of simulated g_j_ time courses in response to a V_j_ step and ramps, using parameters of Cx45 GJs. In these cases, a very small difference between the full model and the REA approximation was observed only at the beginning of a V_j_ step or ramp: the g_j_ curves essentially overlapped elsewhere. An even closer correspondence between the full 4SM and REA approximation was observed when using parameters for Cx43 GJs and evaluating steady-state g_j_-V_j_ relationships (not shown). Overall, the simplified model was almost 90-fold faster than the full 4SM, while providing accuracy well within a 5 percent margin of error for both Cx45 and Cx43 homotypic GJs. Some of our previous models required about 100-fold more CPU time than this simplified model making application of this new model more feasible to account for GJ channels modulation in simulation studies of large clusters of cells.

### Model fitting of heterotypic Cx45/Cx43EGFP GJ channels

To demonstrate the ability of 4SM to model heterotypic GJ channels, we fit junctional currents (I_j_) recorded in Cx45/Cx43EGFP GJs. In these experiments, positive V_j_ steps (as applied to cell-1 that expresses Cx45) caused an increase in g_j_, as can be seen from increase in I_j_ recorded in cell-2 (see Fig. 6A).

**Figure 6.**
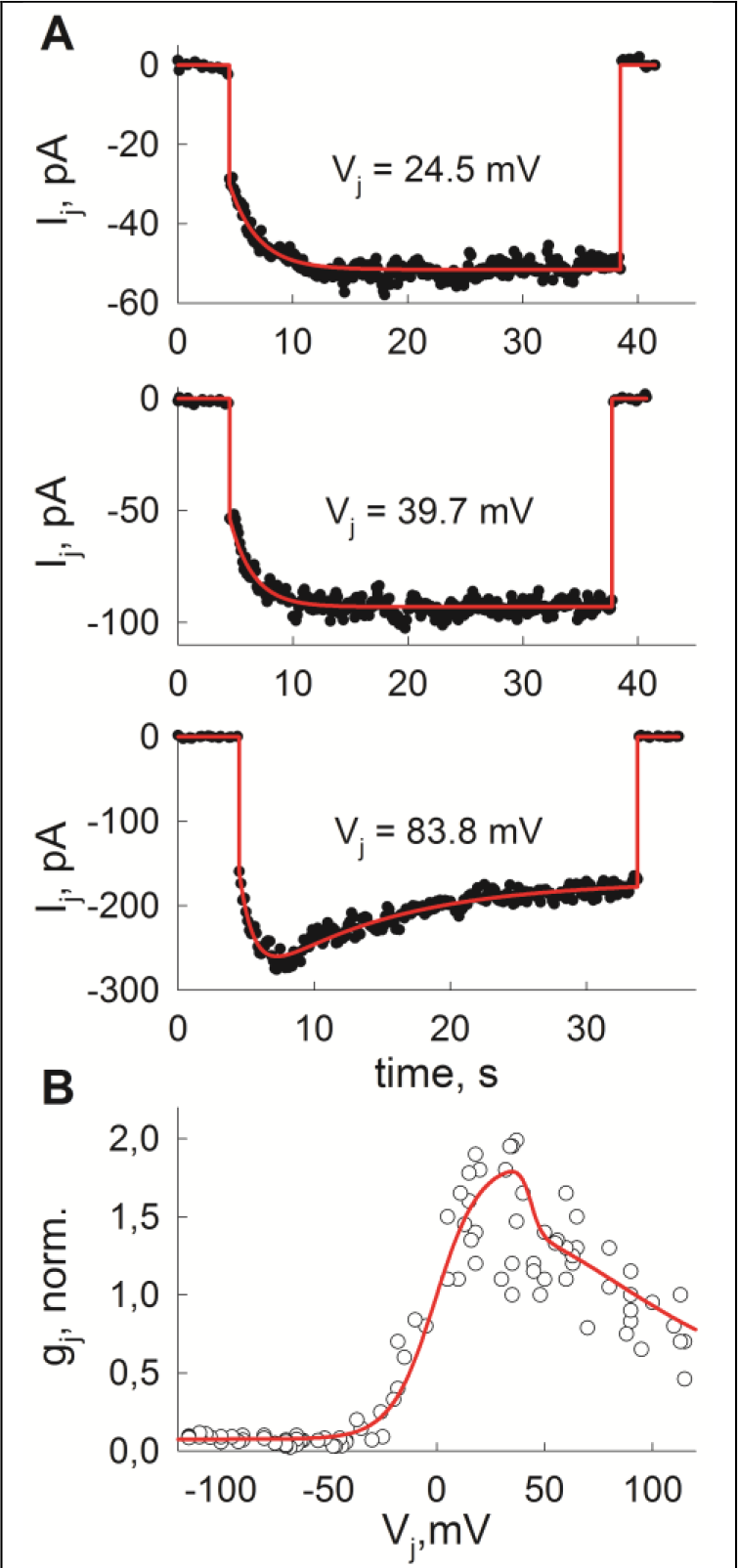
4SM fitting to heterotypic Cx45/Cx43EGFP GJ channels. **(A)** Kinetics of junctional currents (I_j_), caused by V_j_ steps positive on the Cx45 side. Black circles show experimentally recorded values, whereas solid red curves were obtained from 4SM. **(B)** Experimentally observed (white circles) and theoretical (red curve) steady-state g_j_-V_j_ values.

Model fitting of heterotypic GJs using 4SM is more complicated than for homotypic GJs, due to an increase in the number of model parameters (10-12 for heterotypic cases, as compared to only 5 for homotypic cases). Not only does this complication increase computation time during global optimization, but it also makes model fitting less reliable, because very similar g_j_ (or I_j_) time courses can be obtained by different sets of model parameters. For these reasons, we included not only the kinetics of I_j_, but also the steady-state g_j_-V_j_ values into the global optimization procedure. Moreover, to reduce the number of model parameters we did not include rectification of unitary conductance and used the values of the unitary conductances of Cx45 and Cx43EGFP hemichannels deduced from single channel recordings of the corresponding homotypic GJ channels. As reported in [20], these conductance values predict the conductance of the Cx45/Cx43EGFP channel as a simple series connection of Cx45 and Cx43EGFP hemichannels. Both Cx45 and Cx43EGFP hemichannel gating polarities were assigned to be negative.

Fig. 6 demonstrates that 4SM can provide a good fit for both I_j_ kinetics and steady-state g_j_-V_j_ relationships. However, the resulting gating parameters for the Cx45 hemichannel, obtained from fitting data with Cx45 heterotypically paired with Cx43EGFP were significantly different from those obtained when fitting data to homotypic Cx45 GJ channels. The V_0_ value of −9.1875 mV was more than 50 percent lower and the ratio γ_*o*_;/γ_*c*_ = 0.039 was almost 5-fold lower for the Cx45 hemichannel in the heterotypic case compared to the homotypic case. In fact, the model did not satisfactorily fit the data for heterotypic Cx45/Cx43EGFP GJ channels when we used the gating parameters obtained from fits to data from homotypic Cx45 channels using global optimization, even when Cx43-EGFP parameters were allowed to vary and were not constrained to values obtained from homotypic Cx43 GJs. Similar discrepancies in V_j_ gating model parameters between Cxs in homotypic and heterotypic GJ configurations were observed in [10]. In the aforementioned study, the authors raised the possibility that such differences in model parameters reflect changes in hemichannel V_j_ sensitivities due to docking interactions. Additional complications such apparent changes in V_j_ sensitivity caused by open channel rectification were not taken into consideration. We examine the influence of rectification of unitary conductance below.

### Evaluation of rectification of unitary conductance on instantaneous and steady-state g_j_-V_j_ relationship of heterotypic GJs

Experimental data has shown that the conductances of both open and residual states of GJ channels can depend on V_j_ [23–25]. In general, rectification of unitary conductance is a property of heterotypic GJs, because the asymmetry in structure can result in an asymmetric distribution of charged residues in the GJ channel pore that impact ionic flux. Rectification of unitary conductance would have the effect of changing the distribution of V_j_ across each hemichannel as a function of voltage. To explore the influence of such rectification on g_j_-V_j_ relationships, we compared simulation results of models using fixed gating parameters for the hemichannels with and without rectification of unitary conductance (Fig. 7). We compared the results on two types of heterotypic GJs – one consisting of hemichannels with the same (negative) gating polarity (left column, Fig. 7), and another with opposite gating polarities (right column in Fig. 7). GJs with hemichannels that close for opposite polarities would only exhibit reductions in g_j_ for one polarity of V_j_ because both hemichannels, which are docked in a head-to-head fashion, are oriented in the same direction with respect to the field generated by V_j_. To simulate rectification of the unitary conductances of the open and closed states, we used the following set of model parameters for the two hemichannels (in mV): R_1;open_ = R_1;closed_ = ±200, R_2;open_ = R_2;closed_ = −100. Fig. 7 shows that these parameters result in moderate rectification of open GJ channel currents (grey solid lines in 7A and B), and somewhat greater rectification of the residual states (see Figs. 7C and D). For comparison, the dashed black line shows an I_j_-V_j_ curve of an ohmic channel at each system state. To combine channel rectification with gating we used 4SM. The unitary conductances of the open and closed states and the V_j_ gating parameters used for simulation are presented in Table 2 (V_0_ was of the same sign as the respective gating polarity). Although these parameters where chosen for demonstration purposes, the general shapes of the steady-state (g_j,ss_) and initial (g_j,init_) g_j_-V_j_ curves resemble those of heterotypic Cx45/Cx43 GJs (both Cxs with negative gating polarities) and Cx26/Cx32 GJs (Cxs with opposite gating polarities).

**Figure 7.**
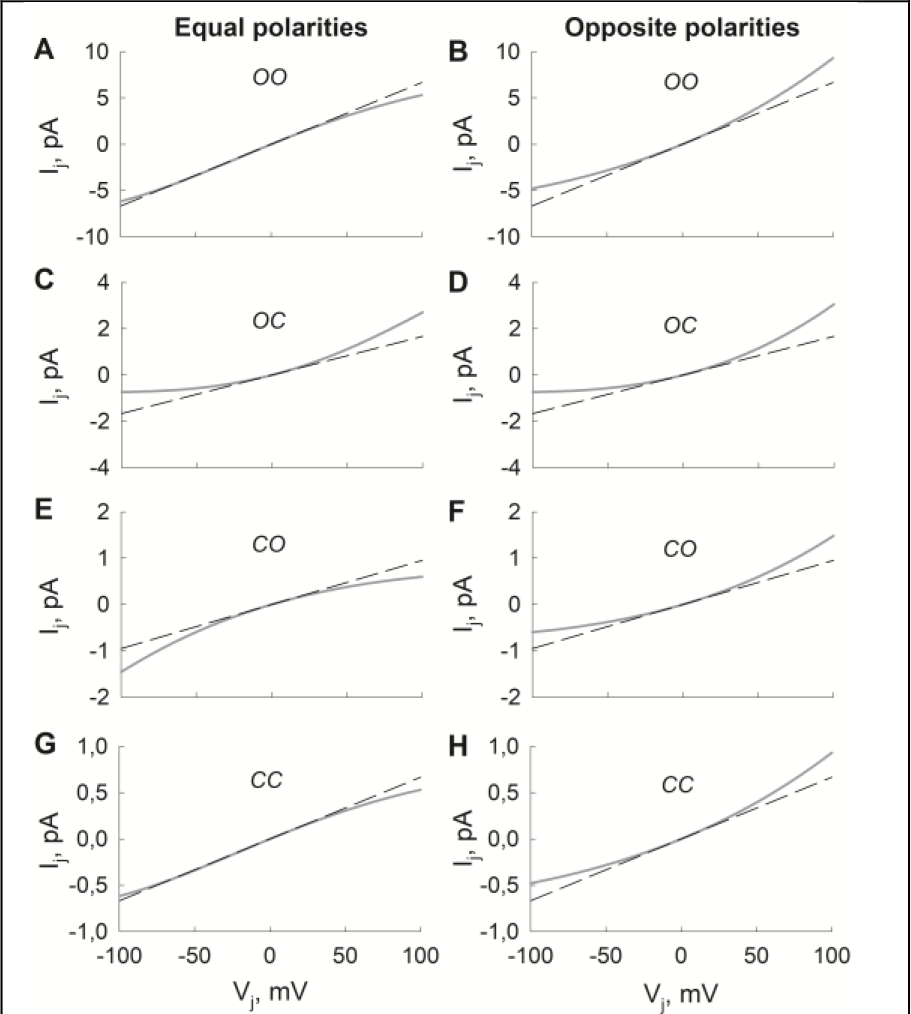
I_j_-V_j_ relationship of rectifying heterotypic chanels at each state of 4SM. Dashed and solid black lines show I_j_-V_j_ curves of nonrectifying and rectifying channels, repectively, associated with each of the indicated system states. Model parameters were the same as presented in Table 2.

**Figure 8.**
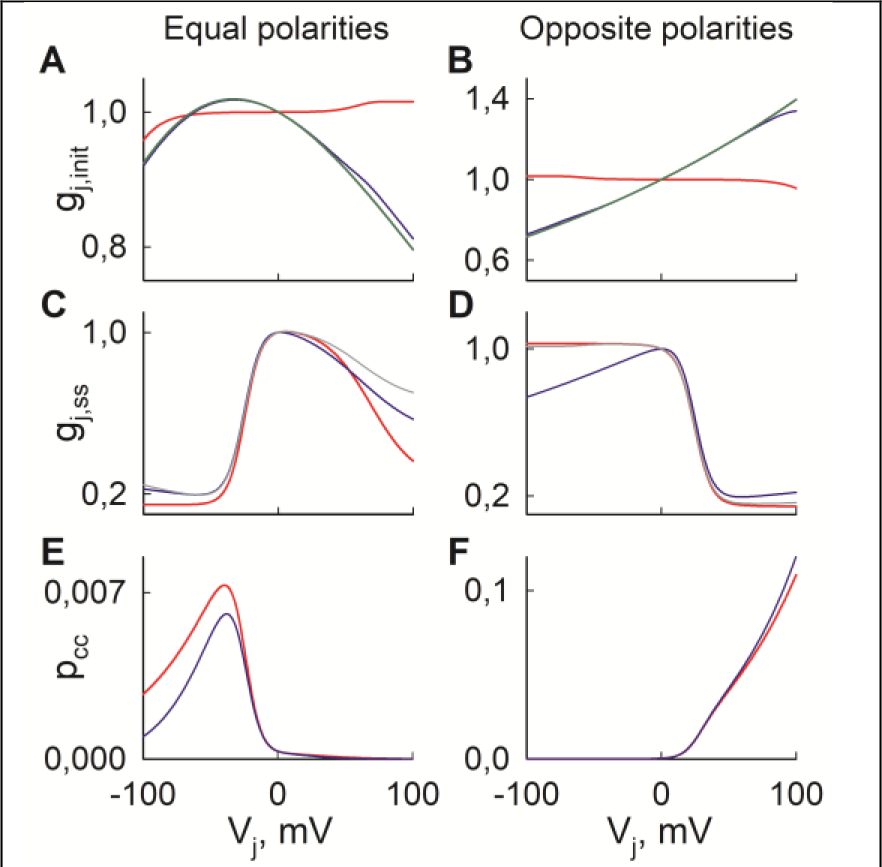
The effect of rectification of unitary conductance in heterotypic GJs. **(A)** and **(B)** Instantaneous g_j_-V_j_ relationships in rectifying (blue line) and non-rectifying (red line) GJs. Green lines show the effect of rectification alone. **(C)** and **(D)** Steady-state g_j_-V_j_ relationship in rectifying (blue line) and non-rectifying (red line) GJs. Grey lines show the ratio of steady-state and instantaneous g_j_ in non-rectifying GJs. **(E)** and **(F)** Steady-state probability of closure of both apposing hemichannels; blue and red lines denote rectifying and non-rectifying GJs, respectively. In all figures, V_j_ is that sensed by the left hemichannel. In the first column, V_j_ gating polarities were negative, i.e., Π_1_=Π_2_=−1; in the second column, Π_1_=1 and Π_2_=−1.

**Table 2.** V_j_ gating parameters used in Fig. 7.

| Hemichannel | $\lambda$, s<sup>-1</sup> | $A_\alpha$, mV<sup>-1</sup> | $A_\beta$, mV<sup>-1</sup> | $V_0$, mV | $\gamma_{open}$, pS | $\gamma_{residual}$, pS |
| --- | --- | --- | --- | --- | --- | --- |
| First | 0.10 | 0.10 | 0.10 | $\pm 10$ | 100 | 10 |
| Second | 0.10 | 0.05 | 0.05 | -40 | 200 | 20 |

It is important to note that V_j_ gating and rectification of channel conductance are interdependent processes, because changes in unitary conductance affect the distribution of V_j_ across each hemichannel. First we examined the initial g_j_-V_j_ curves (Fig. 8A and B), which were obtained by simulating V_j_ steps of 5 ms in duration. To demonstrate the influence of channel rectification, shown are outputs of three different models, which depict: 1) V_j_ gating alone (red line); 2) rectification alone (green line); 3) both V_j_ gating and rectification (blue line). It can be seen that for both types of heterotypic GJs, g_j,init_, the initial conductance measured immediately upon stepping the voltage, is mostly defined by the rectification of the open state. This is expected as g_j,init_ reflects open channel conductance prior to changes in g_j_ that follow as a result of V_j_ gating. Deviation from this correspondence was observed with changes in the V_j_ sensitivity of the hemichannel, in this case the left hemichannel, but was evident only at V_j_s of higher amplitude. For example, when V_0_ of the left hemichannel was changed from ±20 to ±10 mV, the discrepancy between modelling results, which accounted for gating, and the values predicted by rectification alone was ~10 percent at V_j_ = 100 mV. Thus, in many junctions, g_j,init_ would accurately reflect rectification of unitary conductance, especially at low and moderate V_j_s. In cases where V_0_ is severely shifted and/or gating is fast and clamp fidelity is an issue, rectification of the unitary currents should be verified by single channel recordings.

Next we examined the influence of rectification of unitary conductance on the steady-state g_j_-V_j_ relationship (g_j,ss_-V_j_). Because of a noticeable rectification of initial currents in some heterotypic junctions, studies of gating often take the ratio g_j,ss_/g_j,init_ to extract the effects of gating alone. However, this type of transformation. i.e., fitting the g_j,ss_/g_j,init_ curve, does not take into consideration that rectification of unitary conductance changes the distribution of V_j_ across each hemichannel, resulting in a different equilibrium between open and closed states than would be expected in absence of rectification.

Figs. 8C and D shows the results of numerical modeling that combines channel rectification, gating and the effects of their interaction on the distribution of states. Shown are g_j,ss_-V_j_ curves obtained using models with (blue lines) and without (red lines) rectification of unitary conductance; V_j_ gating and rectification parameters were the same as in Fig. 7. The grey curves in Fig. 8C and D show the ratios g_j,ss_/g_j,init_, which were obtained from a model that accounted for both gating and rectification. As can be seen in Fig. 8D, for a heterotypic GJ channel containing hemichannels with opposite gating polarities, g_j,ss_/g_j,init_ values basically overlap with curves without rectification over a large voltage range, and thus reflects the influence of gating alone. However, for a GJ with hemichannels having the same gating polarities (Fig. 8C), g_j,ss_ values with gating and rectification taken into account deviate more from those without rectification (gating alone). Moreover, dividing g_j,ss_ values by g_j,init_ showed even larger deviation from values obtained without rectification. When we performed global optimization for g_j,ss_/g_j,init_ data in Fig. 8C, we obtained the set of model parameters presented in Table 3. Here, the parameter λ was fixed to the same values as in Table 2, because it does not affect g_j,ss_ values. For simplicity, open state conductances were also fixed as in Table 2.

**Table 3.** V_j_ gating parameters, when fitting of non-rectifying GJ model was applied to rectifying GJs

| Hemichannel | $A_\alpha$, mV<sup>-1</sup> | $A_\beta$, mV<sup>-1</sup> | $V_0$, mV | $\gamma_{closed}$, pS |
| --- | --- | --- | --- | --- |
| First | 0.1248 | 0.0778 | -20.50 | 13.67 |
| Second | 0.0173 | 0.1325 | -21.94 | 45.50 |

The results in Table 3 show that model fitting provides a rather different set of V_j_ gating parameters, even though the actual parameter values used were the same as presented in Table 2. For the second hemichannel, estimated V_0_ was ~50 percent lower, while γ_closed_ increased more than twofold, as compared to actual values. This example shows that even moderate rectification of unitary conductance, if not accounted for, can give the false impression that significant changes have occurred in V_j_ gating upon heterotypic docking. Of course, this example does not exclude the possibility that docking significantly affects gating parameters through perturbations in structure, but it demonstrates that an accounting of rectification of unitary conductance is important to assess the extent to which structural perturbations due to docking may have occurred to influence gating.

### Probability of *CC* state

Modelling data for homotypic GJs shows (Fig. 3B) that state *CC* is energetically unfavourable and its steady-state probability, p_cc_, is very low. Our data obtained from simulations of heterotypic GJs showed that a similar condition applies if hemichannels exhibit the same gating polarity. The main difference is that, the peak value for p_cc_ is shifted to a single V_j_ polarity for the heterotypic GJ (see an example in Fig. 8E). Moderate rectification of unitary conductances does not seem to affect p_cc_ significantly.

For heterotypic GJs containing hemichannels of opposite polarities, one could expect to see a substantial value for p_cc_ at a polarity in which both hemichannels tend to close. However, our modelling results show that it is not the case. For example, using the same gating and rectification parameters as in Fig. 7, we found out that p_cc_ only slightly exceeds 0.1 at V_j_=100 mV, while majority of channels still resided in the *CO* state. In this particular example, the more V_j_-sensitive hemichannel closed first, and the majority of V_j_ dropped across it, thus leaving the less V_j_-sensitive hemichannel open; moderate rectification of unitary conductances did not have a significant effect on p_cc_.

These examples show that V_j_ gating of heterotypic GJs can be difficult to predict beforehand, and we believe that mathematical modelling can be a useful additional research tool. For example, model based evaluation of p_cc_ could help to distinguish whether electrophysiological data at a single channel level comes from a channel with both or just a single hemichannel closed.

## Discussion

In this study, we present a 4-state model (4SM) of GJ channel voltage gating. Although it contains less system states than our previous 16-state and 36-state models, this new model describes both kinetic and equilibrium properties of GJ channels, as exemplified by fits of data from Cx45 and Cx43 GJ channels. Moreover, mathematical analyses showed that for homotypic GJs, the model can be further simplified and approximated by a two-state process using rapid equilibrium assumption. This property of the model allows for a very significant reduction in computational time and could be useful in simulation of cell networks, such as in engineered heart slices that represent a confluent multi-layered syncitia of myocytes derived from pluripotent stem cells [26]. In addition, we demonstrated that the model can provide insights into mechanistic behaviour of heterotypic GJs regarding rectification of unitary conductance and probability of closed state.

### Modelling kinetics of junctional conductance

Thus far, most studies of GJ channel gating that have applied mathematical models have only considered steady-state g_j_-V_j_ relationships [27–29], mainly based on Boltzmann functions as presented in [5]. Kinetics have been rarely addressed. In principle, a kinetic model of voltage gating is more useful in that it would describe both kinetic and steady-state changes in g_j_ with V_j_ and provide higher confidence in the correctness of the resulting fitted parameters. It is conceivable that the rate by which g_j_ changes has physiological relevance in cardiac and nervous tissues, where large, rapid and transient changes in transjunctional voltage can occur. Mutations in Cxs have now been linked to 28 distinct disease conditions [30] and in the case of cardiac Cxs, such as Cx40, have been associated with atrial fibrillations [31, 32]. Generally, disease-associated mutations of Cxs are classified as causing loss or altered GJ channel function [30]; the latter is often described as a shift in the steady-state g_j_-V_j_ relationship. However, kinetic modelling would allow one to assess not just the average conductance at all time scales, but also the distribution of channel states underlying the conductances. Such information is likely relevant for a number of physiological processes, as channels residing in residual and open states have different permeability characteristics, thereby differentially influencing electrical and metabolic communication [25]. Kinetic modelling could also be a convenient tool for evaluating voltage gating of Cx channels when using voltage ramp protocols, which provide relevant data on V_j_ gating properties of GJ channels without resorting to the application of the long V_j_ steps needed to obtain steady-state g_j_-V_j_ relationships. The results of model fitting presented here show that 4SM can effectively describe both steady-state and kinetic data. This version of 4SM combines three main features of previously published models: 1) the voltage distribution across apposing hemichannels, connected in series and exhibiting residual conductance as in [11]; 2) rectification of unitary conductance as presented in [8]; 3) a description of opening and closing transition rates as described in [6]. The first two features were included in previously published S16SM [12], but this model was not able to adequately describe gating kinetics. The improved fitting ability of this model is consistent with a more appropriate description of gating transitions based on thermodynamic considerations as presented in [6]. Thus, it is important that this feature be retained upon extension of this model to include additional features of gating, such as the presence of two gating mechanisms in each hemichannel.

### Model fitting assuming a single gating mechanism

Unlike the 16-state and 36-state models (16SM and 36SM), 4SM does not account for the two voltage gating mechanisms known to be present in GJs, slow or loop gating and fast or V_j_ gating [3]. These mechanisms have been shown to be intrinsic to hemichannels. Thus, it was somewhat surprising that the much simpler 4SM model could reproduce the kinetic changes in g_j_ and in some respects, better than the more complex models, 16SM and 36SM, which take both gating mechanisms into account. This result suggests that, at least in the GJ channels we examined here, Cx43 and Cx45, one gating mechanism essentially dominates over a substantial V_j_ range and can largely account for the changes in g_j_. This dominance of one gating mechanism was evident in studies of Cx45/Cx43-EGFP GJ channels, in which the steady-state behaviour of Cx45 hemichannels modelled with one or two gates showed little difference, except at large V_j_s where interactions between two gates becomes evident [20]. To this point, our current model did not reproduce the experimentally observed component of the g_j_-V_j_ curves at high V_j_s. In general, the 4-state model with a kinetic scheme as presented in Fig. 1 could, in principle, result in a two-exponential or even four-exponential decrease in g_j_. However, our theoretical analyses using the REA approximation and model fitting results showed that V_j_ gating properties of GJ channels constrain their behaviour so that the output of a 4-state model is very close to that of a two-state process. In fact, our REA approximation was closer to a full 4-state model at higher V_j_s, providing additional support for the long-held view that a second exponential component, which becomes visible in macroscopic g_j_ recording at V_j_s of higher magnitudes, reveals the influence of the loop gating mechanism. Thus, the current model can be viewed as a “first approximation”, which might be adequate to describe gating kinetics at lower V_j_s or in excitable cell tissues, there V_j_ transients are too short for activation of the loop or slow gating mechanism.

### Estimation and uniqueness of model parameters

Our 4SM model of GJ channel gating does not have a closed form solution. To better estimate model parameters we used global optimization methods, which are often applied in various scientific domains, including systems biology [33]. However, one of the main problems of this approach is the uniqueness of model solutions [34]. Because our model parameters refer to real biophysical properties of Cx channels, it is presumed that a single correct set of parameters should exist for each Cx isoform, although these values might vary depending on species or cell types. In theory, this correct set of model parameters should adequately reproduce g_j_ values measured under any applied V_j_ protocol, with the caveats of experimental variability taken into consideration. Moreover, most global optimization techniques are based on metaheuristic methods for an efficient, but usually random search of the parameter space. If the number of model parameters is high, model fitting of electrophysiological data using global optimization becomes an ill-posed problem. That is, more than one or in some cases practically an infinite number of model parameters can fit a given data equally well. For this reason alone, it is advantageous to start with a simpler model, even though it may lack some recognized mechanistic components. Thus, the presented 4-state model, even though it does not include both gating mechanisms, might offer a good compromise between model applicability and adequacy. For example, our numerical model fitting experiments showed, that global optimization techniques can provide a unique set of (five) model parameters for homotypic GJs using electrophysiological recordings from at least four different V_j_ step protocols or just a single V_j_ ramp protocol. Overall, for homotypic GJs parameter estimation took from 2 to 10 minutes of CPU time (depending on the global optimization method) using a standard desktop computer. For comparison, estimation of 11 different parameters for heterotypic Cx45/Cx43EGFP GJ channels required almost an hour of CPU time. Thus, inclusion of two gating mechanisms, which would require at least doubling of the number of model parameters, would complicate reliable parameter estimation, especially for heterotypic GJ channels. However, for homotypic GJ channels inclusion of the loop gating mechanism into the model should not be much more complicated than application of 4SM to heterotypic GJ channels. The number of model parameters would be, more or less, the same with the possible addition of a variable describing the series conductances associated with the fast and loop gating mechanisms. Thus, the reliability of parameter estimation could be increased using simultaneous fits to multiple data sets obtained using different types of V_j_ protocols. Another useful application would be to correlate electrophysiological recordings at macroscopic and single channel levels, which should increase the chances of finding a correct set of unique model parameters.

### Future considerations

The model parameters in 4SM describe V_j_ gating properties of GJ hemichannels, which typically are inferred from the behaviours of the corresponding GJ channels. However, recordings from undocked hemichannels are possible and provide an opportunity to assess the extent to which docking affects hemichannels incorporated into GJs. Currently, modelling undocked hemichannels would not require an increase in the number of parameters as both gating mechanisms can be assessed in 4SM.

As mentioned previously, an obvious extension of the model would be to include fast and loop gating mechanisms to model GJs. This would require consideration of the kinetic scheme, and as indicated above, makes it more difficult to obtain a unique model solution, as was the case for our previous 36-state model.

Another useful extension would be a model for simulating V_j_ gating at a single channel level. Our current implementation of the model considers averaged behaviours of single channels, and is suitable to describe only macroscopic changes in g_j_. Nevertheless, we believe that the proposed kinetic scheme and the same thermodynamic considerations could be adapted to describe the probabilistic behaviour at the single channel level.

## Author Contribution

M.S. and V.K.V. designed the research. M.S. performed mathematical and computational modeling. T.K., K.M., L.K. and L.G. performed culturing of cells and electrophysiological experiments. T.K. and K.M. collected and analysed electrophysiological data. M.S. and V.K.V. wrote the manuscript with contributions from all authors.

## >Acknowledgement

This research was supported by the Research, Development and Innovation Fund of Kaunas University of Technology (project grant No. PP22/182) and the Science Foundation of Lithuanian University of Health Sciences.

